# Green Synthesis of Silver Nanoparticles Using Sudanese *Candida parapsilosis*: A Sustainable Approach to Combat Antimicrobial Resistance

**DOI:** 10.1101/2025.01.21.634070

**Authors:** Nesreen A. A. Ibrahim, Humodi A Saeed, Samar M. Saeed, Osama Mohamed, Sabah A. E. Ibrahim, Sofia B. Mohamed

## Abstract

**Background:** Antimicrobial resistance (AMR) is a major global health threat, particularly in Sudan, where the overuse and misuse of antibiotics have led to the emergence and spread of multidrug-resistant (MDR) pathogens. Traditional methods to address AMR often fail to provide sustainable solutions. Nanotechnology offers promising alternatives, with silver nanoparticles (AgNPs) demonstrating broad-spectrum antimicrobial properties. This study aims to develop an eco-friendly synthesis of AgNPs using *Candida parapsilosis* isolated from Sudanese soil, leveraging untapped fungal biodiversity to combat AMR.

**Results:** The *Candida parapsilosis*-mediated synthesis of AgNPs was successfully characterized using UV-Vis spectroscopy, X-ray diffraction (XRD), and high-resolution transmission electron microscopy (HRTEM), confirming the formation of well-defined nanoparticles. The biosynthesized AgNPs exhibited potent antimicrobial activity against Gram-positive and Gram-negative MDR pathogens. Medium concentrations of AgNPs demonstrated optimal activity, with inhibition zones up to 29 mm for *Pseudomonas aeruginosa* (ATCC27853). MIC and MBC assays revealed AgNPs’ bactericidal efficacy, particularly against *Escherichia coli* and *Klebsiella pneumoniae* at 0.3125 mg/mL. Synergistic effects with antibiotics, such as ceftazidime and colistin, significantly enhanced antimicrobial activity, with fold increases up to 9.46. AgNPs disrupted bacterial membranes, as evidenced by increased permeability and leakage of nucleic acids.

**Conclusions:** This study presents a novel and sustainable approach to combating AMR by utilizing Sudanese fungal strains for the green synthesis of AgNPs. The findings highlight the potential of AgNPs as an effective antibacterial agent, particularly in combination with conventional antibiotics, to combat multidrug-resistant pathogens. This research not only offers a cost-effective and environmentally friendly solution to AMR but also underscores the significant potential of integrating microbiology and nanotechnology to address global health challenges. The results could pave the way for future applications in both public health and environmental sustainability.

## 1. Introduction

Antimicrobial resistance (AMR) is a critical global health crisis, leading to increased morbidity and mortality worldwide. In Sudan, AMR has emerged as an urgent issue, contributing to an estimated 4.95 million deaths in 2019 due to drug-resistant infections. Multidrug-resistant (MDR) pathogens are prevalent in both hospital and community settings, posing a severe public health challenge. The routine misuse and overuse of antibiotics exacerbate this problem, increasing the prevalence of resistant bacterial strains [1]. This alarming scenario highlights the urgent need for effective alternatives to combat life-threatening bacterial pathogens. Nanotechnology has shown immense potential in addressing AMR. Silver nanoparticles (AgNPs), in particular, have been extensively studied for their remarkable antibacterial, antifungal, and antiviral properties [2]. AgNPs demonstrate broad-spectrum antimicrobial activity by adhering to bacterial cell walls, penetrating into cells, disrupting cellular structures, inducing reactive oxygen species, and altering signal transduction mechanisms [3]. The mechanism through which AgNPs exert their antimicrobial effects primarily involves their interaction with bacterial cell membranes. Upon contact, AgNPs bind to the negatively charged components of the cell membrane, causing physical damage and destabilizing the membrane structure [4]. This disruption leads to increased membrane permeability, leakage of cellular contents, and ultimately cell death. Additionally, AgNPs generate reactive oxygen species (ROS), which further exacerbate membrane damage, DNA fragmentation, and inhibition of cellular respiration [5]. AgNPs also affect bacterial metabolism and enzymatic activities by binding to thiol groups, thereby disrupting vital cellular functions [6]. These properties make AgNPs versatile tools in nanomedicine, capable of inhibiting biofilm formation and acting as drug delivery systems [7].The synthesis of nanoparticles has traditionally relied on chemical and physical methods, which are often energy-intensive and environmentally unfriendly. In recent years, biogenic synthesis has gained attention as a sustainable and cost-effective alternative. Organisms such as bacteria, fungi, and plants can mediate nanoparticle synthesis by producing metabolites that act as natural reducing and stabilizing agents. Fungi, in particular, are considered efficient “nano-factories” due to their ability to produce large quantities of enzymes and proteins, which enable rapid and sustainable nanoparticle synthesis [8]. Moreover, fungi-mediated synthesis allows precise control over the size and morphology of nanoparticles [9]. Globally, fungi such as *Fusarium*, *Aspergillus*, *Trichoderma*, *Verticillium*, *Rhizopus*, and *Penicillium* have been extensively studied for AgNP production [10] .Reports on the biosynthesis of silver nanoparticles (AgNPs) using single-celled yeasts are still limited. However, several yeast species have been investigated for this purpose, including *Saccharomyces boulardii*, *S. cerevisiae*, *Candida albicans*, *Candida utilis*, *Candida lusitaniae*, and yeast strain MKY3. Yeasts are preferred for the extracellular biosynthesis of AgNPs due to their superior tolerance to metals, bioaccumulation properties, and potential for large-scale production, economic viability, and ease of downstream processing. Moreover, yeast produce significant amounts of proteins and enzymes that serve as reducing and stabilizing agents, making them ideal candidates for AgNP synthesis [8]. The nanoparticles have demonstrated significant antimicrobial activity against MDR strains of *Staphylococcus*, *Pseudomonas*, *E. coli*, and *Acinetobacter* [11]. Additionally, biogenic AgNPs have proven effective at low concentrations, with minimal toxicity and negligible environmental impact [12]. Despite this global progress, the potential of Sudanese soil fungi for AgNP synthesis remains largely untapped. Sudan’s diverse ecological landscape likely harbors unique fungal species with the potential to produce bioactive nanoparticles. This study explores the extracellular synthesis of AgNPs using *Candida parapsilosis* isolated from Sudanese soil. The antibacterial activity of these nanoparticles is evaluated, including their synergistic effects when combined with commonly used antibiotics such as ampicillin, cefepime, and vancomycin. This work represents the first comprehensive effort to utilize Sudanese fungi for AgNP synthesis and its application in combating MDR bacteria. By integrating microbiology and nanotechnology, the study offers a green and sustainable approach to addressing the global AMR crisis. The findings underscore the potential of fungal-based AgNPs as a promising alternative in nanomedicine, contributing to both public health and environmental sustainability.

## 2. Material and Methods

### 2.1 Isolation and Identification of Fungi

Soil samples were collected in 2021 from various plant zones in Khartoum State, at four distinct depths, including both surface and sub-surface layers, using standard soil sampling methods (REF). For fungal isolation, 10 grams of soil was suspended in 90 mL of sterile 0.9% NaCl solution and mixed vigorously using a magnetic stirrer for 20–30 minutes to create a uniform suspension. Serial dilutions (up to 10⁻⁵) were performed, and 0.1 mL from each dilution was inoculated onto Potato Dextrose Agar (PDA) plates. The plates were incubated at 25°C for 48–72 hours. Subsequently, fungal strains were cultured on Sabouraud Dextrose Agar (SDA) at 27°C for 1–5 days, depending on the growth rate [13].

### 2.2 Molecular Identification Using 18S rRNA Gene

DNA extraction was carried out using the CTAB method. Fungal material was suspended in a lysis buffer containing 200 mmol/L Tris-HCl (pH 8.0), 0.5% SDS, 250 mmol/L NaCl, and 25 mmol/L EDTA and homogenized using a conical grinder for molds. The suspension was incubated at 100°C for 15 minutes, followed by the addition of 150 µL of 3.0 mol/L sodium acetate and cooling at -20°C for 10 minutes. After centrifugation at 10,000 × g for 5 minutes, the supernatant was extracted sequentially with phenol-chloroform-isoamyl alcohol (25:24:1) and chloroform. DNA was precipitated with isopropanol at -20°C, washed with 70% ethanol, dried, and dissolved in 50 µL of ultrapure water. The internal transcribed spacer (ITS) region was amplified using primers ITS1 (5′-TCCGTAGGTGAACCTGCGG-3′) and ITS4 (5′-TCCTCCGCTTATTGATATGC-3′). PCR conditions included: initial denaturation at 94°C for 2 minutes; 35 cycles of denaturation at 94°C for 30 seconds, annealing at 56°C for 10 seconds, and extension at 72°C for 30 seconds; followed by a final extension at 72°C for 2 minutes [13].

### 2.3 Sequencing and Analysis of the Amplified DNA

PCR products were analyzed by electrophoresis on a 2% (w/v) agarose gel stained with ethidium bromide. Each sample contained 5 µL of PCR product mixed with 1 µL of gel loading dye. A molecular marker (50– 2000 bp) was used to determine the fragment size, and *Saccharomyces cerevisiae* (RPMCC 9763) was used as a positive control (Pryce et al., 2003). The amplified DNA was purified using QIAquick Gel Extraction Kits (Qiagen). Sequencing was performed using the BigDye™ Terminator v3.1 Cycle Sequencing Kit (Applied Biosystems). Sequences were analyzed using the BLAST algorithm in the NCBI database for phylogenetic identification. A phylogenetic tree was constructed using the Neighbor-Joining method in MEGA v11.0 software [14].

### 2.4 Biosynthesis of Silver Nanoparticles (AgNPs)

The *Candida parapsilosis* isolate was evaluated for its ability to biosynthesize silver nanoparticles (AgNPs). The biomass of the isolate was prepared by culturing it aerobically in Potato Dextrose Broth (PDB), formulated using 250 g of potato and 20 g of dextrose per liter of distilled water. The culture suspension was incubated on an orbital shaker at 25 ± 2 °C with agitation at 120 rpm for 96 hours. Following incubation, the biomass was harvested through filtration and thoroughly washed with distilled water to remove any residual medium components. Subsequently, 10 grams of the washed biomass were suspended in 100 mL of sterilized double-distilled water and incubated for 48 hours at 25 ± 2 °C in a 250 mL Erlenmeyer flask with continuous agitation at 120 rpm. The resulting suspension was filtered using Whatman filter paper No. 1 (GE Healthcare, Buckinghamshire, UK) to obtain the cell-free filtrate. This filtrate was then treated with a 1 mM solution of silver nitrate (AgNO₃) in an Erlenmeyer flask and incubated at room temperature in a dark environment to facilitate nanoparticle synthesis (Devi and Joshi, 2012) [13].

### 2.5 Characterization of Silver Nanoparticles (AgNPs)

The absorption peaks of AgNPs colloidal solutions were analyzed using several techniques. First, visual observation revealed a gradual color change of the cell-free filtrate from fungal isolates to brown upon incubation with a silver nitrate solution under dark conditions, indicating nanoparticle formation. Second, a CARY-100 BIO UV-Vis Spectrophotometer (Varian Inc., Palo Alto, CA, USA) with a resolution of 1 nm was used to record the absorption spectra, confirming the presence of AgNPs. Third, X-ray diffraction (XRD) measurements were conducted using a Philips X-ray generator model PW 3710/31 (Philips, Japan) with a diffractometer featuring an automatic sample changer (PW 1775), a scintillation counter, a Cu-target tube, and a Ni filter, operating at 40 kV and 30 mA, with a scan range of 5–80° (2θ). Finally, Transmission Electron Microscopy (TEM) analysis was performed with a JEOL JSM 100CX TEM instrument (Jeol, Tokyo, Japan) operated at 200 kV to examine the morphology and size of the nanoparticles [13].

### 2.6 Re-identification of Multidrug-Resistant Isolates through Macroscopic, Microscopic, Biochemical, and API-Based Approaches

The re-identification of MDR isolates involved several diagnostic approaches. Microscopic examination was carried out using Gram staining, focusing on Gram reaction, shape, and arrangement. Biochemical tests followed standard procedures for bacterial diagnosis. The API Staph system was employed to identify *Staphylococcus* species, where bacterial suspensions were inoculated into microtubes containing dehydrated substrates, and results were determined based on color changes. Additionally, the API 20E Kit was used for the identification of *Enterobacteriaceae* and *Pseudomonas aeruginosa* through a series of biochemical tests conducted in microtubes with specific substrates [15].

### 2.7 Antimicrobial Susceptibility Test

Antimicrobial susceptibility testing was conducted following standard protocols. McFarland Standard was prepared by mixing barium chloride and sulfuric acid to estimate bacterial concentrations, with absorbance measured at 625 nm to ensure proper turbidity. The modified Kirby-Bauer method was employed for susceptibility testing, where bacterial suspensions were adjusted to match the 0.5 McFarland standard and inoculated onto Muller-Hinton agar plates. After brief drying, antimicrobial disks (including ampicillin, amikacin, vancomycin, ciprofloxacin, and others) were applied and incubated at 37°C for 18-24 hours to assess bacterial resistance.

### 2.8 Antimicrobial Activity of AgNPs

The antimicrobial activity of the synthesized AgNPs was tested against various bacterial strains, including *E. coli* (ATCC 43890), *E. faecalis* (MTCC 2729), *S. aureus* (MTCC 96), *Pseudomonas* (ATCC 27853), and clinical isolates of *K. pneumoniae*, *E. coli*, *A. baumannii*, *Pseudomonas spp.*, *S. typhi*, *Serratia*, *S. aureus*, *E. faecalis*, *L. monocytogenes*, and *Bacillus spp.* from urine, wound, and blood samples collected from hospitals in Khartoum, Sudan. The antimicrobial activity was assessed using the agar well diffusion method. Wells (9 mm in diameter) were created in Mueller-Hinton Agar (MHA) plates, and different volumes (100 μL, 500 μL, and 1000 μL) of AgNP solutions were added to test high, medium, and low concentrations. Water extract without mycelia was used as a negative control. The plates were incubated at 37°C for 48–72 hours, and inhibition zones were measured. The disk diffusion method was used to evaluate the synergistic effects of AgNPs with antibiotics. A 20 μL solution of AgNPs was added to each antibiotic disk, which was then placed on MHA plates inoculated with test organisms. The plates were incubated at 25°C for 24–48 hours, and the diameters of inhibition zones were measured. The experiment was repeated three times for accuracy and reliability [13].

### 2.9 Minimum Inhibitory Concentration (MIC) and Minimum Bactericidal Concentration (MBC) Determination

MIC and MBC were determined using the broth dilution method (CLSI M07-A8). Serial dilutions of AgNPs were prepared, ranging from 5 mg/mL to 0.156 mg/mL. The bacterial concentration was adjusted to 10⁸ CFU/mL using the 0.5 McFarland standard. These dilutions were tested in BHI broth, and the control sample consisted of broth without inoculation. The tubes were incubated at 37°C for 24 hours, and the MIC was defined as the lowest concentration of AgNPs with no visible bacterial growth. For MBC determination, 50 μL samples from tubes showing no growth were plated on MHA agar and incubated at 37°C for 24 hours. The MBC was the lowest concentration of AgNPs that killed 99.9% of the bacterial population [16].

### 2.10 Synergistic Effects of AgNPs with Resistant Antibiotics

The synergistic effects of AgNPs with commonly used antibiotics were evaluated using the disk diffusion method. A 20 μL solution of AgNPs was applied to standard antibiotic disks, which were placed on MHA plates inoculated with test organisms. The plates were incubated at 25°C for 24–48 hours, and the diameters of the inhibition zones were measured. The experiment was repeated three times for consistency [13].

### 2.11 Membrane Integrity and Permeability In Vivo Activity

The integrity of the bacterial cell membrane was assessed by measuring the release of DNA and RNA, which absorb at 260 nm, using specific concentrations of AgNPs. The bacterial solution was treated with AgNPs, and permeability changes were measured by assessing the absorbance at OD420 over time using O-nitrophenyl β-D-galactopyranoside (ONPG) as an indicator [17].

### 2.12 Statistical Analysis

Data were analyzed using statistical package for social science software (SPSS version .20) Independent T test were performed to quantitative and quantitative variables. A P.value of< 0.05 was considered as significant for all statistical tests in the present study.

## 3. Result

### 3.1 Fungi isolation and identification

Two strains of *Candida parapsilosis*, designated CSS1 (accession number PQ796639) and CSS2 (accession number PQ796640), were isolated from soil samples collected in Khartoum, Sudan. A phylogenetic tree was constructed to confirm their taxonomic identification (Figure 1). These fungal isolates were subsequently utilized for the biosynthesis of silver nanoparticles (AgNPs) using silver nitrate (AgNO₃) as the precursor, and their antibacterial activity was evaluated.

**Fig 1:**
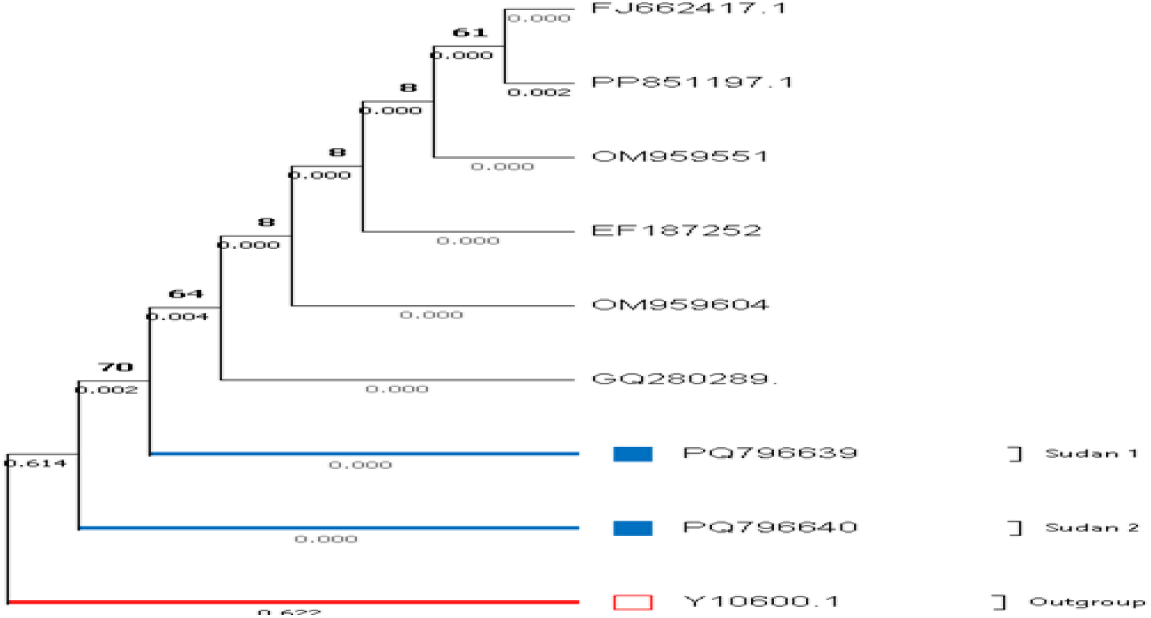
Phylogenetic Tree Confirming the Identification of *Candida parapsilosis* Strains CSS1 (PQ796639) and CSS2 (PQ796640)

### 3.2. Biosynthesis of AgNPs

The synthesis of AgNPs resulted in distinct solution colors corresponding to three different concentrations: low concentration (L), medium concentration (M), and high concentration (H) (Figure 2).

**Fig. 2.**
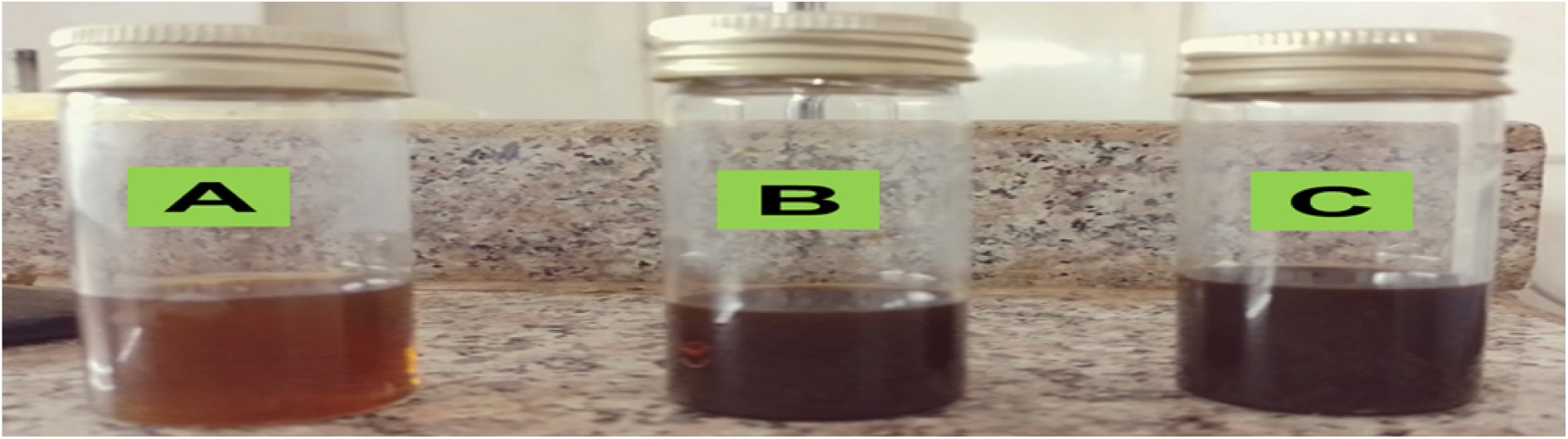
The biosynthesis results of AgNPs demonstrated varying solution colors depending on the concentration: (A) AgNPs at low concentration (L), (B) AgNPs at medium concentration (M), and (C) AgNPs at high concentration (H)

### 3.3 Characterization of AgNPs

The analysis of the synthesized silver nanoparticles (AgNPs) using various methods revealed distinct findings. As shown in (Figure 3), the UV-Vis absorption spectra of AgNPs synthesized under different conditions exhibit surface plasmon resonance (SPR) peaks within the range of 400–450 nm, confirming the formation of AgNPs. Variations in peak intensity, position, and width indicate differences in nanoparticle size, uniformity, and aggregation. A sharper, more intense peak (blue curve) suggests the presence of smaller, more uniform nanoparticles, while broader or less intense peaks (red and black curves) indicate larger particles or increased aggregation. These results highlight how synthesis conditions affect the optical and structural properties of the AgNPs.

**Fig. 3.**
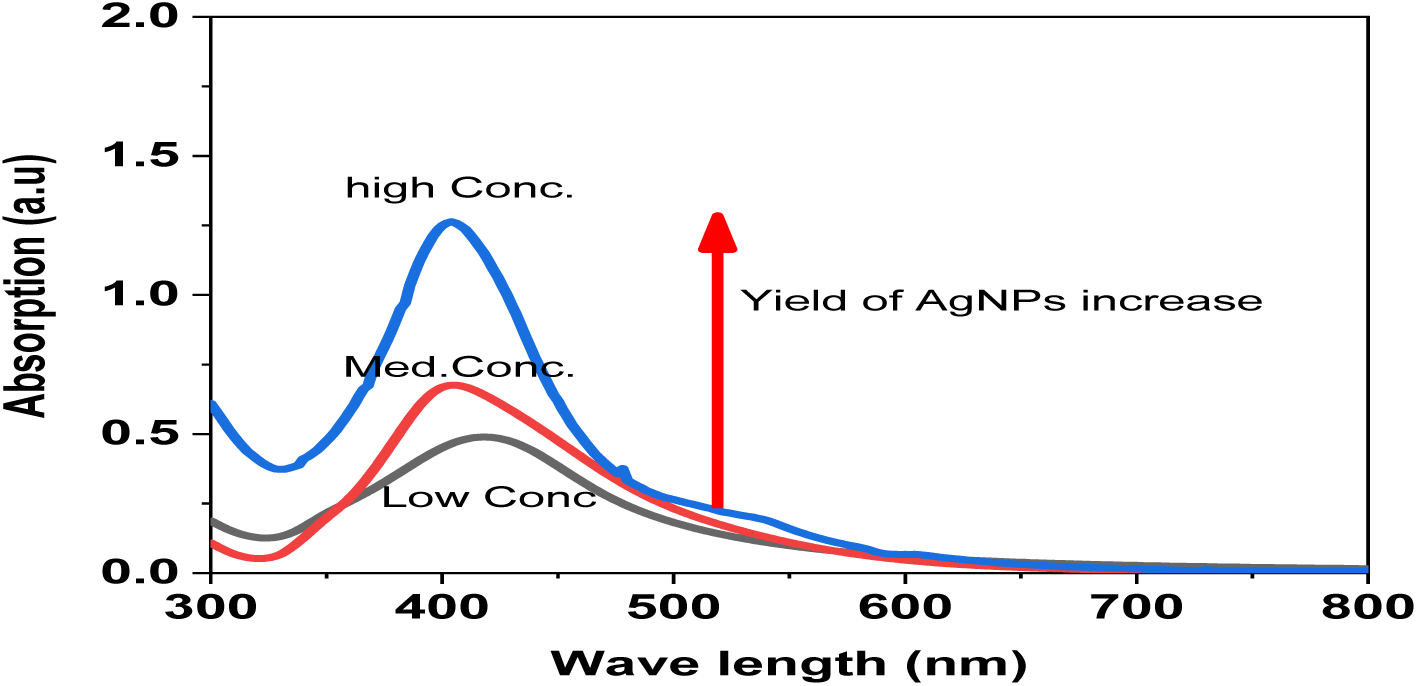
UV-visible absorption spectra of biosynthesized AgNPs at varying concentrations following exposure to silver nitrate solution.

The X-ray diffraction (XRD) analysis, presented in (Figure 4), showed distinct diffraction peaks at 2θ angles of 37.8°, 45.9°, 65.1°, and 77.01°, corresponding to the crystallographic planes (111), (200), (220), and (311), confirming the crystalline nature of the AgNPs.

**Fig. 4.**
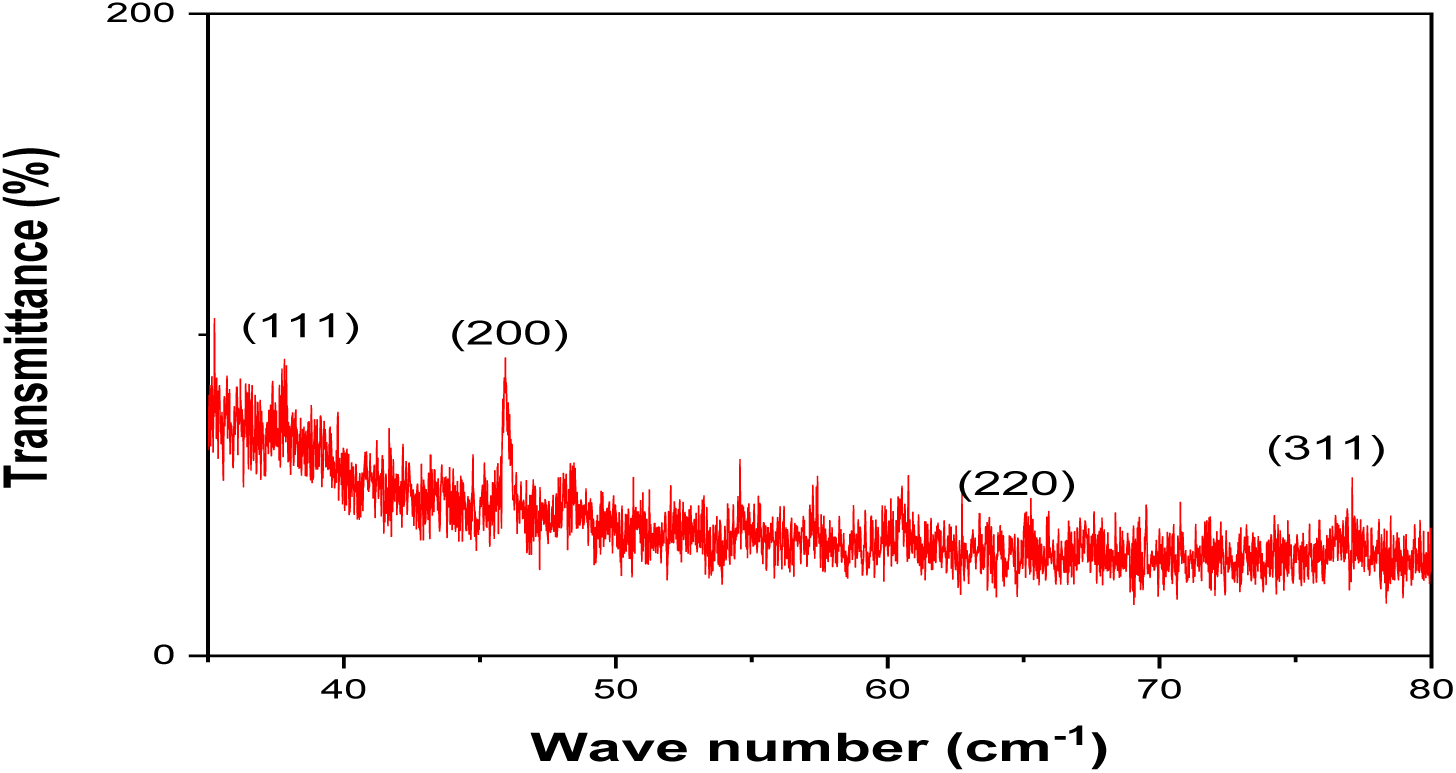
XRD diffraction pattern of biosynthesized silver nanoparticles (AgNPs).

The TEM images and histograms in (Figure 5) illustrate the impact of precursor material concentration on the characteristics of silver nanoparticles (AgNPs). At low concentrations, the nanoparticles appear smaller with less aggregation, while the histogram indicates moderate size uniformity (Fig 5a,b). Medium concentrations result in more uniform and well-dispersed nanoparticles with minimal aggregation, as reflected by the narrower size distribution in the histogram (Fig 5c,d). At high concentrations, significant aggregation is evident, leading to larger, irregular clusters, with the histogram showing a wider size distribution and reduced control over particle size (Fig 5e,f). These findings highlight that medium precursor concentrations are optimal for achieving a balance between particle size uniformity and minimal aggregation. High concentrations, on the other hand, lead to excessive aggregation, which may negatively affect the functional performance of the nanoparticles in applications such as catalysis, antimicrobial activity, or optical technologies. The results emphasize the importance of carefully controlling precursor concentrations during synthesis, particularly when scaling up production, to ensure consistent nanoparticle characteristics.

**Fig. 5.**
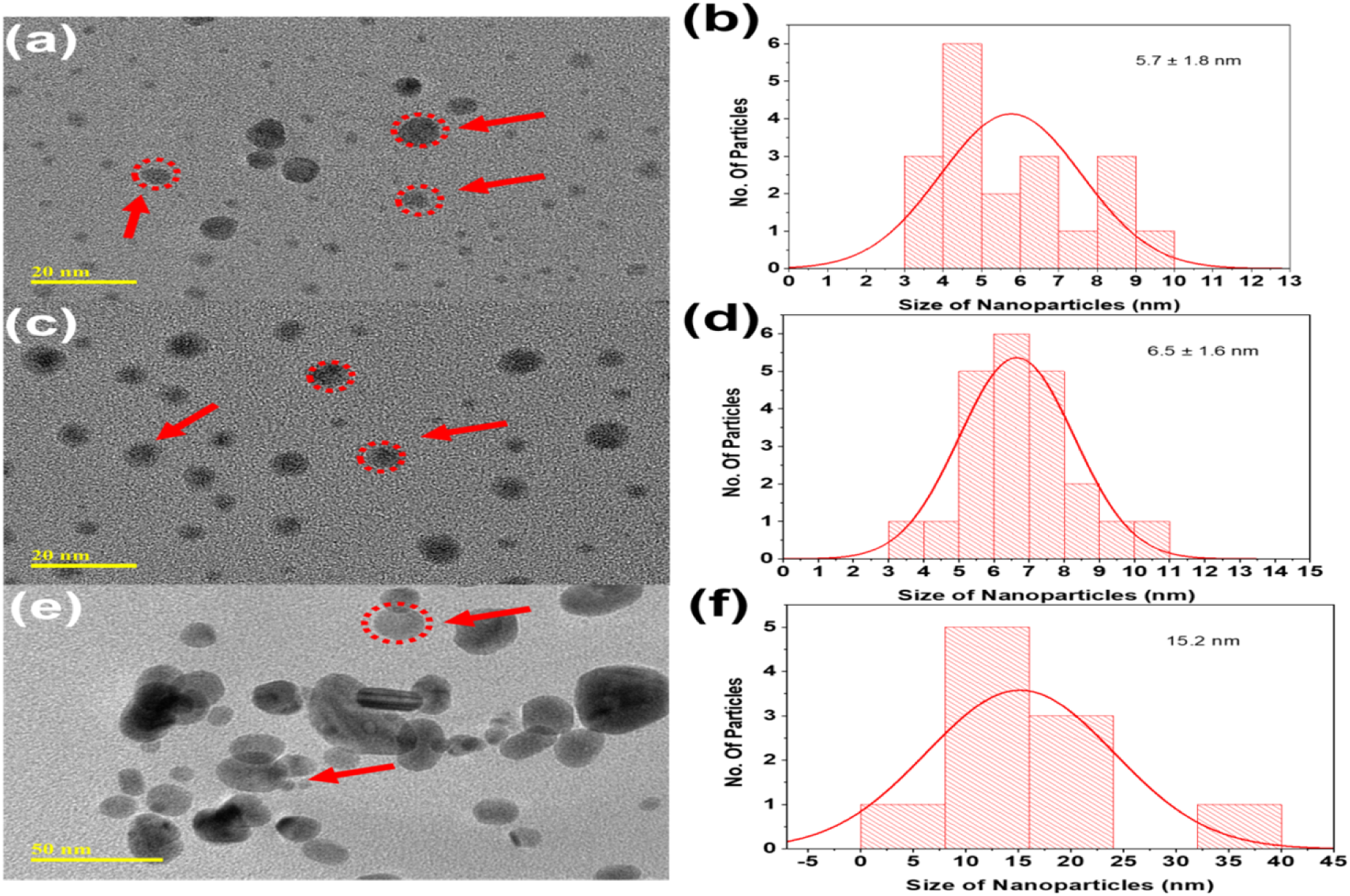
TEM images and corresponding histograms of biosynthesized AgNPs: (a) AgNPs at low concentration with (b) their particle size distribution histogram, (c) AgNPs at medium concentration with (d) their histogram, and (e) AgNPs at high concentration with (f) their histogram.

(Figure 6) presents High-Resolution Transmission Electron Microscopy (HRTEM) analysis, confirming the successful biosynthesis of AgNPs with the following key findings: Morphology (100 nm scale): The AgNPs are predominantly spherical or nearly spherical, with some aggregation, and their sizes are within the nanometer range (Fig 6A). Higher Magnification (50 nm scale): At higher magnification, the spherical shape of the nanoparticles is confirmed, offering a detailed visualization of individual particles (Fig 6 B). Crystalline Structure (5 nm scale): Lattice fringes observed at high magnification reveal the crystalline nature of the AgNPs, with a lattice spacing of 0.24 nm corresponding to the (111) plane of metallic silver (Fig 6 C). Selected Area Electron Diffraction (SAED) Pattern: The bright concentric rings in the diffraction pattern indicate a polycrystalline structure, confirming the metallic silver composition of the nanoparticles (Fig 6 D).Line Profile Analysis: Numerical analysis of lattice fringes shows interplanar distances (e.g., 0.2359 nm and 0.2588 nm), matching theoretical values for silver’s crystal planes (Fig 6 E). These findings validate that the synthesized AgNPs are highly crystalline and hold significant potential for various applications.

**Fig. 6.**
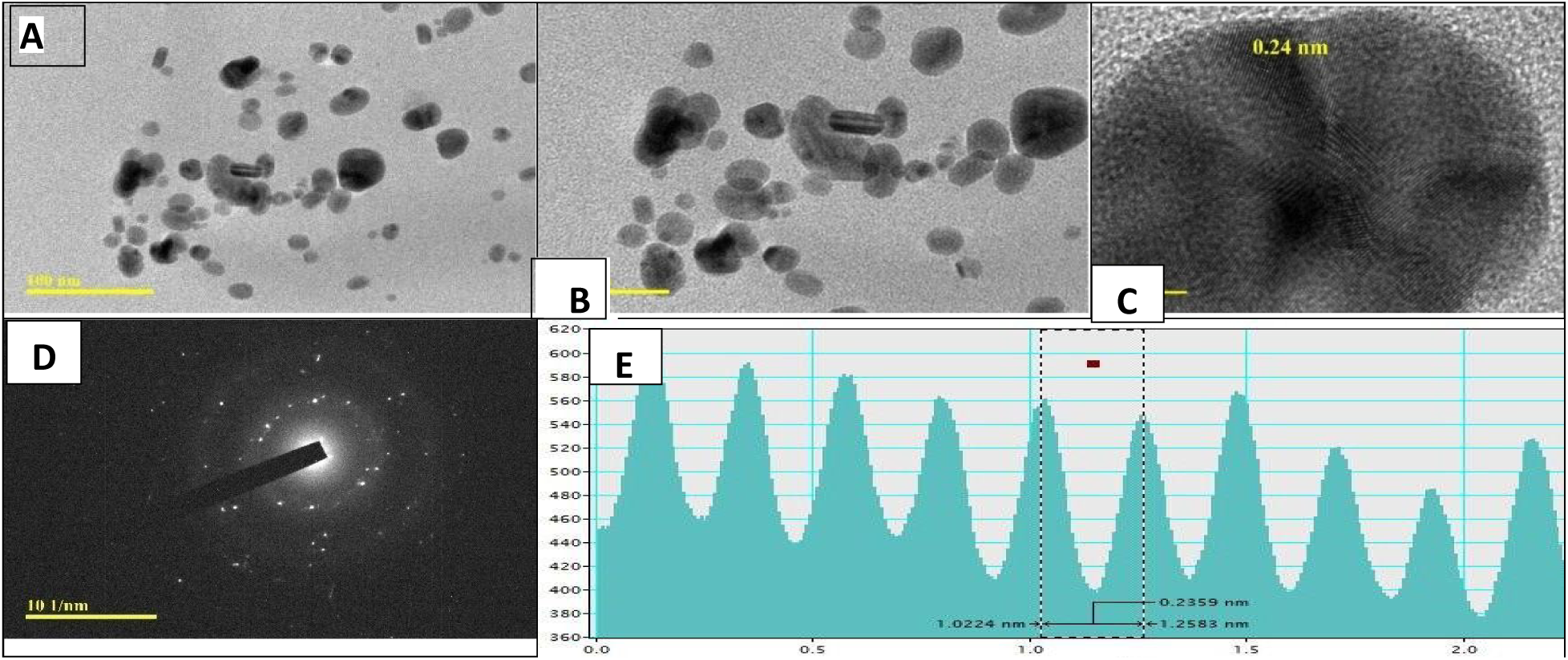
HRTEM analysis of biosynthesized AgNPs: (A) HRTEM image with a scale bar of 100 nm, (B) magnified image with a 50 nm scale bar, (C) high-resolution image with a 5 nm scale bar, (D) selected area electron diffraction (SAED) pattern, and (E) line profile showing interplanar lattice fringes.

### 3.4 Antibiotic Susceptibility of Isolated Bacteria

The study assessed the antibiotic susceptibility of ten pathogenic bacteria against twelve common antibiotics to determine their resistance profiles. *E. coli* showed complete resistance to Ampicillin and Vancomycin but remained sensitive to Chloramphenicol and Imipenem. *Klebsiella pneumoniae* exhibited broad resistance, particularly to Ampicillin and Amikacin (S Table 1). *Pseudomonas aeruginosa* and *Acinetobacter baumannii* were resistant to multiple antibiotics but sensitive to Piperacillin, Imipenem, and Trimethoprim Sulphate (S Table 2).

*Salmonella typhi* showed resistance to Bacitracin and Cephalexin but was sensitive to Amikacin and Imipenem. *Serratia marcescens* was resistant to Bacitracin but sensitive to Amoxicillin Clavulanate (S Table 3). *Staphylococcus aureus* was resistant to Colistin and Tetracycline, yet sensitive to Imipenem. *Enterococcus faecalis* was fully resistant to Amikacin and Ampicillin (S Table 4). *Listeria monocytogenes* showed resistance to Nitrofurantoin but remained sensitive to several antibiotics. *Bacillus cereus* was resistant to Aztronam and Nitrofurantoin but sensitive to others (S Table 5).

These findings highlight the variability in antibiotic resistance across bacterial species, emphasizing the need for personalized antibiotic treatment to effectively combat resistance.

### 3.5 Antimicrobial Activity of AgNPs

Silver nanoparticles (AgNPs) exhibited antibacterial activity against ATCC reference bacteria, with medium concentrations demonstrating the highest inhibition zones across all strains. For *E. coli* (ATCC43890), inhibition zones ranged from 22 mm (high concentration) to 24 mm (medium concentration). *E. faecalis* (MTCC2729) and *S. aureus* (MTCC96) showed the greatest activity at medium concentration (28 mm), while high concentrations resulted in reduced efficacy. *P. aeruginosa* (ATCC27853) showed optimal inhibition at medium concentration (29 mm), with a decline at high concentration (25 mm). The reduced activity at higher concentrations may be due to nanoparticle aggregation or diffusion interference. Low concentrations showed moderate but consistent antibacterial activity. These results highlight medium concentration as the most effective dose, emphasizing the need for optimized nanoparticle usage to maximize antimicrobial efficacy (Fig S1) (Table 1).

**Table 1.** Inhibitory Effect of Biosynthesized AgNPs at High, Medium, and Low Concentrations Against ATCC Reference Bacterial Strains.

| ATCC | Low (mm) | Medium (mm) | High (mm) |
| --- | --- | --- | --- |
| *E. coli- ATCC43890* | 23 | 24 | 22 |
| *E.faecles - MTCC2729* | 21 | 28 | 22 |
| *S. aureus- MTCC96* | 22 | 28 | 24 |
| *P. aeruginosa-ATCC27853* | 24 | 29 | 25 |
| **Mean± SD** | 22.5±1.3 | 27.3± 2.2 | 23.2± 1.5 |

Silver nanoparticles (AgNPs) demonstrated antibacterial activity against both Gram-positive and Gram-negative bacteria at high (500 mg/ml), medium (250 mg/ml), and low (100 mg/ml) concentrations. The highest inhibition zones were observed at high concentrations (mean: 21.0 ± 2.79 mm), with *S. typhi* (25 mm) and *E. coli* (24 mm) being the most susceptible. Medium concentrations showed comparable efficacy (mean: 20.1 ± 2.89 mm), particularly against *P. aeruginosa* and *A. baumannii*. Low concentrations had the lowest activity (mean: 17.7 ± 3.34 mm), with minimal inhibition observed in *S. marcescens* (10 mm). Gram-negative bacteria exhibited higher susceptibility compared to Gram-positive species (Fig S2) (Table 2).

**Table 2.** Inhibitory Effect of Biosynthesized AgNPs at High, Medium, and Low Concentrations Against Various Gram-Positive and Gram-Negative Bacterial Species.

| Bacteria species | Concentration |  |  |
| --- | --- | --- | --- |
|  | High (500mg/ml)/100ml | Medium (250mg/ml)/50ml | Low (100mg/ml)/25ml |
| *E. coli* | 24 | 23 | 17 |
| *K. pneumonia* | 20 | 18 | 17 |
| *P. aeruginosa* | 22 | 24 | 20 |
| *A. baumannii* | 23 | 23 | 21 |
| *S. marcescens* | 18 | 18 | 10 |
| *S. typhi* | 25 | 22 | 21 |
| *S. aureus* | 22 | 20 | 20 |
| *E. faecalis* | 19 | 18 | 18 |
| *L. monocytogenes* | 16 | 15 | 15 |
| *B. cereuse* | 21 | 20 | 18 |
| **Mean± SD** | 21.0±2.79 | 20.1±2.89 | 17.70±3.34 |

### 3.6 Antimicrobial Activity of Silver Nanoparticles (AgNPs) via Tube Dilution Method for MIC and MBC

Silver nanoparticles (AgNPs) exhibited bactericidal activity against all tested Gram-negative bacteria, with MBC/MIC ratios ≤1. *E. coli* and *K. pneumoniae* showed the highest sensitivity, with both MIC and MBC at 0.3125 mg/mL. *P. aeruginosa* and *A. baumannii* demonstrated moderate sensitivity, with MIC at 1.25 mg/mL and MBC at 0.625 mg/mL. Similarly, *S. typhi* and *S. marcescens* had MIC of 0.625 mg/mL and MBC of 0.3125 mg/mL. These findings highlight the potent antibacterial efficacy of AgNPs against Gram-negative bacteria, particularly for *E. coli* and *K. pneumoniae.* (Table 3).

**Table3.** Minimum Inhibitory Concentration (MIC) and Minimum Bactericidal Concentration (MBC) of AgNPs (mg/mL) Against Gram-Negative Bacteria.

| Compounds | AgNPs |  |  |
| --- | --- | --- | --- |
|  | MBC | MIC | Ratio MBC/MIC |
| *E. coli* | 0.3125 | 0.3125 | 1 |
| *K. pneumonia* | 0.3125 | 0.3125 | 1 |
| *P. aeruginosa* | 0.625 | 1.25 | 0.5 |
| *A.baumannii* | 0.625 | 1.25 | 0.5 |
| *S. typhi* | 0.3125 | 0.625 | 0.5 |
| *S. marcescens* | 0.3125 | 0.625 | 0.5 |

Silver nanoparticles (AgNPs) exhibited variable antibacterial activity against Gram-positive bacteria. *S. aureus* showed a MBC/MIC ratio of 1, indicating bactericidal activity. *L. monocytogenes* and *B. cereus* were the most susceptible, with MBC/MIC ratios of 0.5, reflecting strong bactericidal effects. *E. faecalis* displayed a higher MBC/MIC ratio of 2, suggesting a bacteriostatic effect at lower concentrations. These findings demonstrate the effectiveness of AgNPs, particularly against *B. cereus* and *L. monocytogenes*, as potent antibacterial agents (Table 4).

**Table 4.** Minimum Inhibitory Concentration (MIC) and Minimum Bactericidal Concentration (MBC) of AgNPs (mg/mL) Against.

| Compounds | AgNPs |  |  |
| --- | --- | --- | --- |
|  | MBC | MIC | Ratio MBC/MIC |
| *S.aureaus* | 0.625 | 0.625 | 1 |
| *E. faeclies* | 1.25 | 0.625 | 2 |
| *L. monocytogenes* | 0.3125 | 0.625 | 0.5 |
| *B.cereuse* | 0.1562 | 0.3215 | 0.5 |

### 3.7 Synergistic Activity of AgNPs Coated on Antibiotics Against Resistant Bacteria

The synergistic effect of AgNPs when combined with common antibiotics was assessed for several resistant bacterial strains. (S Table 6) shows that the addition of AgNPs significantly enhanced the antimicrobial efficacy of various antibiotics. The data indicated fold increases in antibacterial activity, with the effect being strain- and antibiotic-specific. Among the antibiotics tested, *Ceftazidime* demonstrated the most substantial enhancement, with a 5.11-fold increase (p < 0.001) in its activity against *Acinetobacter*. AgNPs also showed significant synergistic effects with *Ceftazidime* (8.19-fold, p < 0.001) and *Cefepime* (5.93-fold, p < 0.01) against *Bacillus* species. Moderate improvements were observed with *Ampicillin* (1.02), *Erythromycin* (1.28), and Nitrofuran (1.80) against *E. coli*, while Colistin exhibited a remarkable 2.85-fold increase (p < 0.05). For *Enterococcus*, the fold increase was most pronounced with Colistin (9.46-fold, p < 0.001) and Ceftazidime (3.84-fold, p < 0.01). The combination of AgNPs with Ampicillin (2.70-fold) and Ceftazidime (2.33-fold) effectively increased susceptibility in *Klebsiella* species. AgNPs greatly enhanced the effectiveness of Tetracycline (11.5-fold, p < 0.001) and Colistin (2.84-fold, p < 0.05) against *Listeria*. The combination of AgNPs with Ceftazidime (4.04-fold) and Gentamicin (2.63-fold) showed enhanced efficacy against *Pseudomonas*, while for *S. aureus*, AgNPs enhanced Ceftazidime (3.74-fold) and Colistin (4.48-fold) effectiveness. The synergistic effects observed for *Serratia* (Ceftazidime: 7.63-fold; Colistin: 7.87-fold) and *Salmonella* (Ceftazidime: 3.64-fold; Colistin: 4.84-fold) highlight the capacity of AgNPs to enhance antibiotic activity against resistant strains.

### 3.4. Confirmatory the antibacterial activity of AgNPs

The antibacterial effect of AgNPs on bacterial membrane integrity was assessed using three line graphs comparing treated and untreated bacterial cells. The data showed significant alterations in membrane integrity following AgNP treatment. Across all graphs, the red lines (post-treatment) demonstrated a marked increase in the measured parameter, such as membrane permeability or damage, compared to the gray lines (control), indicating the disruptive effect of AgNPs on bacterial membranes. In the (Fig 7 a), membrane integrity steadily increased after AgNP treatment, suggesting a continuous disruption of the bacterial membranes. In (Fig 7 b), a sharp rise in membrane damage was noted immediately following AgNP treatment, followed by a slower yet sustained increase, while the control group showed minimal changes. The (Fig 7 c) exhibited a sharp increase in membrane damage post-treatment, which peaked at a midpoint and then plateaued or slightly declined, suggesting a strain-specific or concentration-dependent effect. These findings confirm the antibacterial efficacy of AgNPs in disrupting bacterial membranes in a manner that is dependent on both concentration and bacterial strain. The differences in response among bacterial strains highlight the selective activity of AgNPs, suggesting their potential for targeting specific bacterial pathogens.

**Fig.7.**
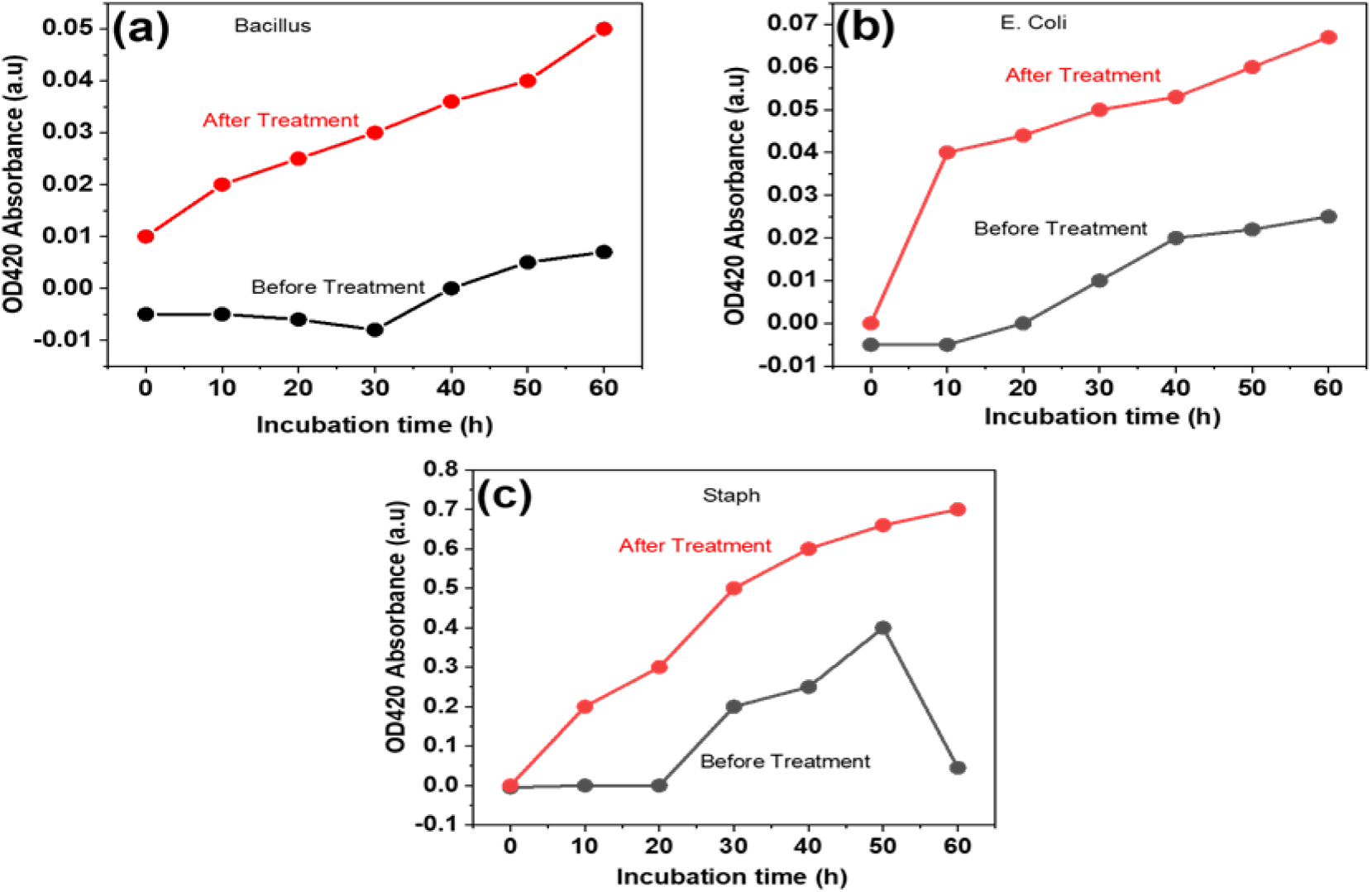
Line chart depicting the membrane permeability of representative bacterial isolates before and after treatment with green-synthesized AgNPs: (a) *Bacillus*, (b) *E. coli*, and (c) *S. aureus*.

## **4-** Discussion

The biosynthesis of silver nanoparticles (AgNPs) using fungal isolates from Sudanese soil represents a novel and eco-friendly approach to nanotechnology, with significant implications for combating antimicrobial resistance (AMR). Unlike conventional methods of nanoparticle synthesis, which rely on toxic chemicals and energy-intensive processes. The biosynthesis of AgNPs using single-celled yeasts is still relatively unexplored, with only a few yeast species reported in the literature such as *Candida albicans* and *Candida lusitaniae* [8]. This study demonstrates a sustainable alternative by harnessing the natural enzymatic capabilities of *Candida parapsilosis*.

Characterization of synthesized AgNPs through UV-Vis spectroscopy, XRD, and TEM confirmed their nanoscale size, crystalline nature, and uniform spherical morphology. These findings are consistent with prior report using *Fusarium oxysporum*, which yielded similar nanoparticle characteristics [18].

In this study, medium concentrations of silver nitrate (AgNO3) produced smaller AgNPs (5.7 to 6.5 nm) with a narrow size distribution, while higher concentrations resulted in larger particles with broader size distributions. The surface plasmon resonance band observed at 419 nm confirmed the particle size. The smaller AgNPs exhibited greater antibacterial activity compared to other studies [19]. For instance, a study in Saudi Arabia using desert soil fungi produced AgNPs with sizes ranging from 10 to 16 nm [19], while this study achieved larger inhibition zones against *Escherichia coli* and *Staphylococcus aureus*.

The synthesized AgNPs exhibited remarkable antibacterial activity against Gram-positive and Gram-negative multidrug-resistant (MDR) bacteria, including *E. coli*, *S. aureus*, and *P. aeruginosa*. Medium concentrations of AgNPs consistently provided optimal inhibition zones, attributed to their smaller size and effective interaction with bacterial cells. These results align with findings from earlier studies on fungi-mediated AgNPs, which highlighted their ability to inhibit bacterial growth by disrupting membrane integrity [20].

The antibacterial activity demonstrated in this study exceeds many previously reported findings. For example, the inhibition zones for *E. coli* and *S. aureus* using the fungi-mediated AgNPs in this study were 20 mm and 23 mm, respectively. These results are markedly higher than those reported in earlier studies, where inhibition zones were 12 mm for E. coli and 8.3 mm for *S. aureus* (18). Similarly, research utilizing *Fusarium sp*. documented inhibition zones of 16 mm for *E. coli* and 13 mm for *S. aureus* [13], both of which are lower than the values observed in our study.

The synergistic effects of AgNPs combined with antibiotics underscore the innovative nature of this research. Previous studies reported modest increases in inhibition zones, with fold enhancements of up to 1.89 when combining AgNPs with antibiotics [13]. In contrast, the current study achieved substantially higher fold increases, including a 9.46-fold enhancement with colistin against *Enterococcus faecalis* and a 7.87-fold enhancement with ceftazidime against *Serratia*. These results highlight the superior efficacy of the AgNPs synthesized in this study, likely attributed to their smaller size and distinctive surface properties.

Mechanistically, the antibacterial action of AgNPs involves disruption of bacterial membrane integrity and permeability, as evidenced by increased release of nucleic acids and reduced membrane stability post-treatment. These findings are consistent with previous studies that demonstrated similar effects, including depolarization and destabilization of bacterial membranes by AgNPs [21]. The observed differences in inhibition zones, minimum inhibitory concentrations (MICs), and minimum bactericidal concentrations (MBCs) across bacterial strains highlight the selectivity and efficiency of the synthesized nanoparticles.

The exceptional antibacterial efficacy of the synthesized AgNPs can be attributed to their distinctive spherical morphology, high crystallinity, and small size. These characteristics enable stronger interactions with bacterial cell walls, resulting in membrane destabilization, increased permeability, and leakage of intracellular contents. Membrane integrity assays conducted in this study revealed substantial damage to bacterial membranes, aligning with previous findings that reported similar mechanisms of action for AgNPs [19].

Interestingly, the ability of AgNPs to disrupt bacterial membranes was more pronounced in Gram-negative bacteria such as *E. coli,* likely due to their thinner peptidoglycan layer, which facilitates nanoparticle penetration. These findings align with previous studies [22, 23] but demonstrate enhanced efficacy, likely due to the optimized biosynthesis process used in this research.

Overall, the study underscores the potential of fungi-mediated biosynthesis of AgNPs as a sustainable and effective strategy for addressing the global challenge of antimicrobial resistance. By utilizing *Candida parapsilosis* isolates from underexplored regions such as Sudan, this research contributes to advancing nanotechnology while providing practical solutions for combating MDR pathogens. These findings reinforce the broader applicability of green-synthesized AgNPs in medical, environmental, and industrial applications.

## 5 Conclusion

This study successfully demonstrated the novel eco-friendly synthesis of silver nanoparticles (AgNPs) using *Candida parapsilosis* strains isolated from Sudanese soil, emphasizing the potential of local fungal biodiversity for sustainable nanotechnology applications. The synthesized AgNPs exhibited unique spherical morphology and high crystallinity, contributing to their remarkable antibacterial efficacy against both Gram-positive and Gram-negative bacteria, including multidrug-resistant (MDR) strains. The findings revealed the ability of AgNPs to disrupt bacterial membrane integrity and enhance the effectiveness of conventional antibiotics, highlighting their potential to combat antimicrobial resistance (AMR).This synergistic interaction not only reduces the required antibiotic doses but also mitigates the spread of resistant pathogens. By integrating green nanotechnology and microbiology, the study provides a sustainable approach to addressing AMR, enhancing public health outcomes, and contributing to environmental preservation in in Sudan and globally. Future work should focus on scaling up production, evaluating in vivo applications, and ensuring safety for broader clinical use. This pioneering effort underscores the value of leveraging indigenous microbial resources to tackle global health challenges and paves the way for further advancements in fungal-based nanoparticle synthesis and application.

## Declarations

### Ethics approval and consent to participate

This study received ethical approval from the National University Research Ethic Committee (NU-REC), National University-Sudan (Approval Number: [NU-RECG121]). The ethical review was conducted in accordance with the National Statement on Ethical Conduct in Human Research (2007), which align with the ethical principles outlined in the Belmont Report and the Declaration of Helsinki.

Written informed consent was obtained from all adult participants prior to their inclusion in the study. Participant privacy and confidentiality were safeguarded throughout the research, and personal identifiers were removed from all datasets.

This study does not involve minors, vulnerable populations, or clinical interventions and is not classified as a clinical trial. Consequently, clinical trial registration is not applicable.

### Consent for publication

Not applicable

### Availability of data and materials

The nucleotide sequences obtained from the fungal isolates have been deposited in the GenBank database and assigned unique accession numbers (PQ796639 and PQ796640).

### Authors’ contributions

HA supervised the study, while SM and SB served as co-supervisors. NA was responsible for sample collection, isolation and identification of fungi and bacteria, synthesis of AgNPs, evaluating the synergistic activity of AgNPs, and confirming their antibacterial properties. NA and OM carried out the molecular techniques and sequencing. SA performed the statistical analysis, and SB conducted the bioinformatics analysis. The first draft of the manuscript was written by HA, SM, SB, and NA, with SB writing the final draft. All authors read and approved the final manuscript.

## Acknowledgements

Not applicable

